# Physico-chemical properties of co-formulation of insulin with pramlintide

**DOI:** 10.1101/227363

**Authors:** Dayana Cabral da Silva, Luís Maurício T. R. Lima

**Author notes:** Corresponding Author: Luis Maurício T. R. Lima – School of Pharmacy, Federal University of Rio de Janeiro – UFRJ, CCS, Bss24, Ilha do Fundão, 21941-590, Rio de Janeiro, RJ, Brazil. Twitter: @MauTrambaioli.

## Abstract

Since the discovery of amylin its use has been discouraged by the inadequacy of the protocol involving multiple injections in addition to insulin. While a combined fixed-dose formulation is thus highly desirable, it has long been limited due to incompatibility as historically documented. We have investigated the compatibility of regular and fast-acting insulin analogues (Aspart, Asp^B28^, and LisPro, Lys^B28^Pro^B29^) with the amylin analogue pramlintide. Insulin interacts with pramlintide, forming heterodimers as probed by electrospray ionization - ion mobility spectrometry-mass spectrometry. While their interaction is likely to delay the amyloid aggregation of pramlintide in phosphate-buffered solution pH 7.0, they do not prevent aggregation at this condition. At acidic sodium acetate solution pH 5.0, combination of pramlintide and the fast-acting insulin analogues become stable against amyloid aggregation. The co-formulated product at high concentration of both pramlintide (600 μg/mL,150 μM) and LisPro insulin (50 IU/mL, 300 μM) showed also stability against amyloid aggregation. These data indicate a potential for the development of a co-formulation of fast-acting LisPro insulin with pramlintide, which could bring benefits for the combined therapy.

**Abbreviations:**

IAPP,: islet amyloid polypeptide;
ESI-IMS-MS,: Electrospray Ionization–Ion Mobility Spectrometry–Mass Spectrometry.

## INTRODUCTION

Amylin (also known as islet amyloid polypeptide – IAPP) is a peptide hormone co-secreted with insulin by the pancreatic β-cell (1,2). Amylin display several physiologic and pharmacologic roles such as control of appetite, gastric emptying, restoring leptin sensitivity, regulation of glucagon secretion, among others (3–6). Amylin is deficient in individuals with type 1 diabetes (T1DM) and advanced stages of type 2 diabetes (T2DM), it is found oversecreted during the subclinical diabetes (7,8) and found aggregated as amyloid fibrils in advanced stages of T2DM (9,10).

Since the discovery of amylin three decades ago (11,12) prompted for an immediate therapeutic interest (patent US 5,367,052 (13)), which was hampered by the limited solubility of human amylin and its propensity for amyloid aggregation. Aggregation of amylin was discovered to be minimized in proline-rich variants (14), which inspired the design of the triple-proline human amylin analogue named pramlintide (Pro^25,28,29^; CAS #151126-32-8; acetate salt CAS #196078-30-5; patent US 5,998,367 (15)). Although proline-rich variants such as murine amylin and pramlintide still holds amylodogenic features (16–18), they show improved physico-chemical properties and similar pharmacologic properties of human amylin (19) resulting in its approval by the FDA in 2004 and entering the US market as first-in-class in 2005 (20). Other approaches have been developed aimed to overcome the stability problem of amylin analogues, such as PEGylation ((21,22) patent US 20,160,331,811) and molecular confinement into liposomes (23) or polymeric particles (21). Given the importance of amylin physiology and pharmacology, others amylin analogues, hybrids and receptors agonists are currently found in pipeline of development (24,25).

Despite the benefits observed with incorporation of pramlintide to the therapeutic scheme, the incompatibility of pramlintide analogues and insulin in a combined formulation (26) requires administration through separated injection (27), becoming a burden for patients due to the multiple daily injections through the day, discouraging its use and becoming a deterministic factor in the lack of adherence to the therapy.

While insulin has long been shown to interact reciprocally with amylin (28,29), further details on the interaction with pramlintide is unknown. Insulin and pramlintide are also likely to show a physico-chemical incompatibility due to the formulation pH: while regular insulin and fast-acting insulin analogues are formulated at neutral solution (pH 7 and higher), pramlintide is formulated at pH 4, resulting in an instability issue in their reciprocal pH (17,30–32). While a prompt precipitation of each individual component may take place when outside the optimal pH range, there is limited information on the structural features of them under these circumstances.

Previous attempts have been made in order to obtain a co-formulation product of insulin with pramlintide. A study showed that extemporaneous combined formulation comprising both regular insulin and pramlintide did not altered their pharmacokinetics profile when compared to the isolated products (33), although this protocol did not addressed issues concerning physicochemical stability. Besides these results, the recommendation for not mixing both compounds holds. Some patent applications have addressed the issue of co-formulation of insulin and amylin analogues (WO 1,999,034,764 = US 6,136,784; WO 2,013,177,565 = US 2,0150,174,209) (34–36), although no successful results was shown concerning the stability of fast-acting insulin with pramlintide.

Here we report the investigation of the physico-chemical interaction and compatibility of pramlintide and a variety of insulin analogues, the regular and the fast-acting analogues Aspart (Asp^B28^) and LisPro (Lis^B28^Pro^B29^), and the morphologic characterization of the combination products. We show their interaction by ESI-IMS-MS, the amyloid aggregation process and the achievement of a particular co-formulation with insulin and pramlintide, with further discussion on the implication for the potential development of a combined formulation comprising both hormones.

## MATERIAL AND METHODS

### Materials

– Carboxy-amidated, C2-C7 dissulfide bond pramlintide (TFA salt, lot #89437, >95% purity) was purchased from Genemed Biotech Inc (CA, USA). The stock peptide solutions were prepared at 10 mg/mL (2.55 mM) in DMSO and stored at −20 °C until use. Insulin products were purchased from local drugstores, stored at 4°C, used within the due date, and aliquot were taken for immediate use. All other reagents were of analytical grade. We used a conversion factor of 1 μIU/mL equivalent to 6.0 pM according to the World Health Organization (WHO) standard of 26,000 IU/g human insulin (37,38).

### Electrospray Ionization–Ion Mobility Spectrometry–Mass Spectrometry (ESI-IMS-MS)

– ESI-IMS-MS measurements were performed in a MALDI-Synapt G1 (Waters Brazil; analytical facility provided by UEMP-UFRJ) high definition quadrupole-travelling wave mass spectrometer (HDMS). Samples were prepared at 50 μM final concentration of each peptide (Pramlintide, Insulin Regular/Aspart/Lispro) in 50 mM ammonium acetate (adjusted with either NH_4_OH or acetic acid) pH 5.0 or pH 7.0, and injected at 2 μL/min, using a positive ESI with a capillary voltage of 2.8 kV and N_2_g at 0.4 bar. Data acquisition was conducted over the range of *m/z* 500 to 4,000 for 10 min. Calibration was performed with phosphoric acid. Other typical instrumental settings are as described previously (16). Data were analyzed using DriftScope 2.4 (39) (Waters Corporation, Brazil).

### Aggregation assay

– kinetic aggregation assays were performed in 96-well flat-bottom black plate (Corning #3915) sealed with transparent film (Duck Brand Crystal Clear Tape, Avon, OH). Experiments were performed at indicated polypeptide concentration from their respective stock solution (either insulin, commercial formulations at 3.45 mg/mL = 300 μM, or pramlintide at 10 mg/mL in DMSO, equivalent to 2.55 mM), at 25 °C, and 50 mM buffer (either Na_2_HPO_4_ or sodium acetate), in a final volume of 200 μL. Measurements were continuously performed in a SpectraMax M5 microplate reader (Molecular Devices) with excitation set at 440 nm and emission at 482 nm, PMT medium, 30 flash/well, and a 475 nm cut-off filter, with 3 min time interval preceded by 3 sec vibration.

### Transmission electron microscopy (TEM)

– The products of the aggregation assays were evaluated for morphology by TEM by depositing 5 μL of the suspension onto a Formvar-coated Cu grids (300 mesh), removed after about 2 min and contrasted with 2% uranyl acetate. TEM images were acquired in a Zeiss 900 transmission electron microscope operating at 80 kV, in the CENABIO-UFRJ facility.

### Co-formulation of LisPro insulin with pramlintide

– We used commercially available LisPro insulin (100 IU/mL) to concentrate in Centricon Ultracel^®^ 3K (cut-off 3000 NMWL; Amicon^®^ Ultra 2mL, Ref: UFC200324 Lot:R4HA81570) by centrifugation at 4,500 x g at 4 °C until reaching a reduction to about 4x the original volume. The LisPro insulin was diluted back to approximately its original concentration using ice-cold 50 mM sodium acetate buffer pH 5.0, and again concentrated. This procedure was repeated 5 times. The final protein concentration after buffer exchanging was determined by the fluorescamine method (40) and was added of pramlintide (from original solution in DMSO at 10 mg/mL). The sample was mixed gently and left overnight at 4 °C for immediate use in amyloid aggregation assays. All attempts to perform the same procedure with the Aspart insulin failed due to aggregation during the concentration step.

## RESULTS

### Insulin:pramlintide interaction by ESI-IMS-MS

– The interaction between insulin analogues and pramlintide and formation of stable oligomers was evaluated by mass spectrometry. Equimolar amount of insulin and pramlintide were combined in 50 mM ammonium acetate buffer and injected into an electrospray ionization – mass spectrometry couplet to ion mobility spectrometry (ESI-IMS-MS). This technique allows the separation of similar, overlapping mass/charge ions by differential mobility in a gas-pressurized chamber, and has been previously used in the characterization of varying oligomeric states in the stepwise association of both murine amylin (16) and pramlintide (17).

Heterodimers of insulin and pramlintide in charged state +5 have been detected for all insulin variants, i.e., regular (**Fig. 1**), Aspart (**Fig. 2**) and LisPro (**Fig. 3**), both at pH 5.0 and pH 7.0, along with monomers and homodimers. While homodimers of insulin could be detected for the regular variant, only minor signal of homodimers was observed for Aspart and LisPro insulin analogues at present assay conditions. This results is indicative of reduced prevalence of such oligomeric state – homodimers - in the fast-acting insulin analogues compared to the regular insulin, a known physico-chemical feature of these fast-acting insulin analogues as previously reported (41). Heterodimers with charged state +4 of pramlintide with these same insulin analogues have also been detected at both pH 7.0 and pH 5.0 (**Fig. 4**). These data confirm the ability for homo- and hetero-functional assembly of these insulin variants and pramlintide, although the outcomes of such interaction remained elusive until presentation of such as described below.

**Figure 1.**
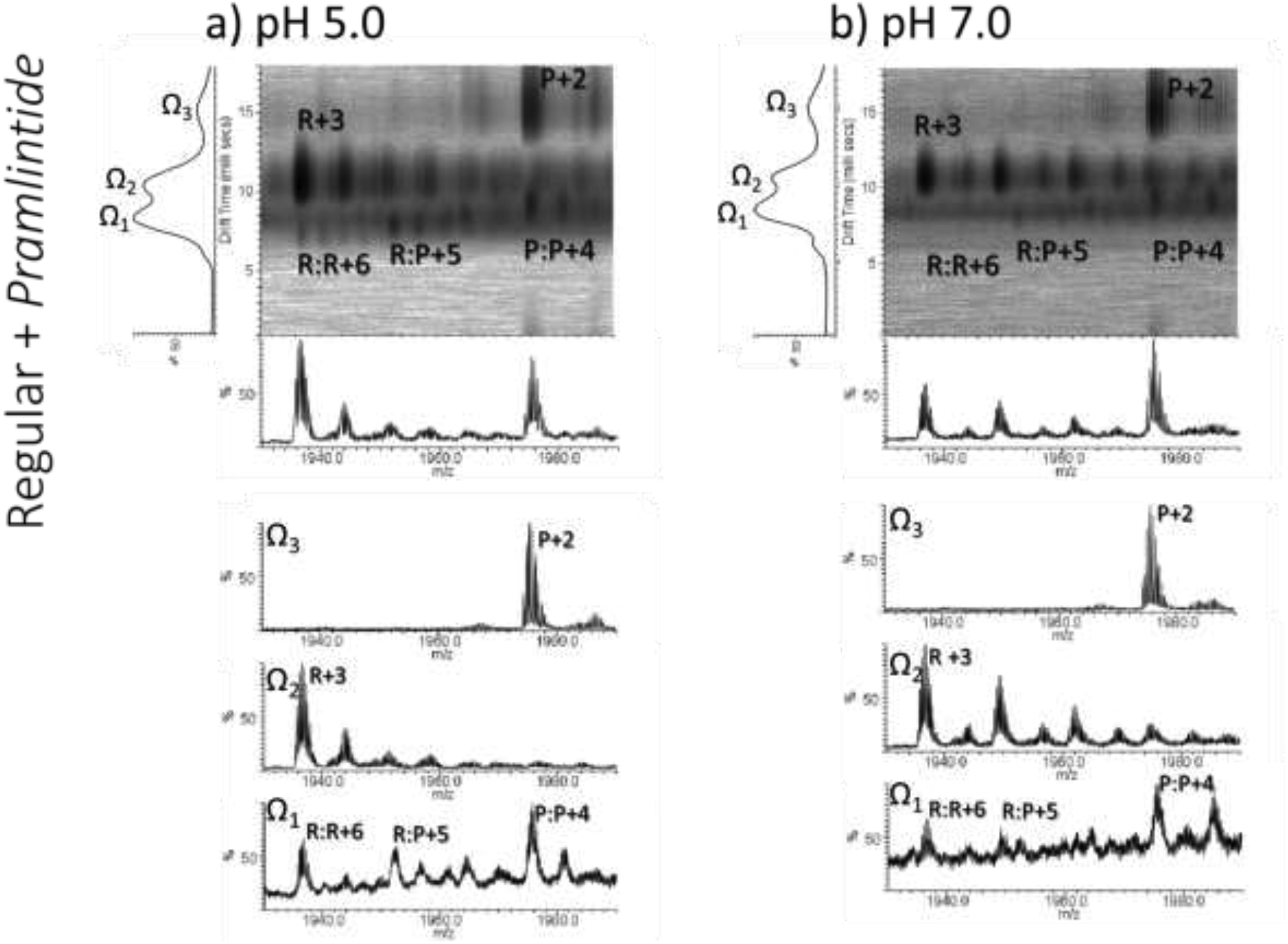
Distribution of oligomeric states of regular insulin and pramlintide accessed by electrospray ionization – ion mobility – mass spectrometry (ESI-IMS-MS). Regular insulin (**R**) and pramlintide (**P**) were combined at 50 μM each in 50 mM ammonium acetate pH 5.0 (a) or pH 7.0 (b) and assayed for ESI-IMS-MS. Monomers are represented by a single letter (either **R** or **P**) and homodimers or hetero-dimers are identified by the combination of their corresponding letters. Their respective charge state, represented by number following the ‘plus’ sign, is shown adjacent to the identification of the oligomeric state. The ion intensity in the drift time (*d*t) axis x *m/z* is plotted in square root scale, and in linear scale for the total ESI-MS spectrum along the whole *d*t axis shown below the *d*t x *m/z* panel. A stripping of the *d*t x *m/z* space comprising specific *d*t regions (Ω_1_, Ω_2_ and Ω_3_) as assigned in the figure are shown for clarity.

**Figure 2.**
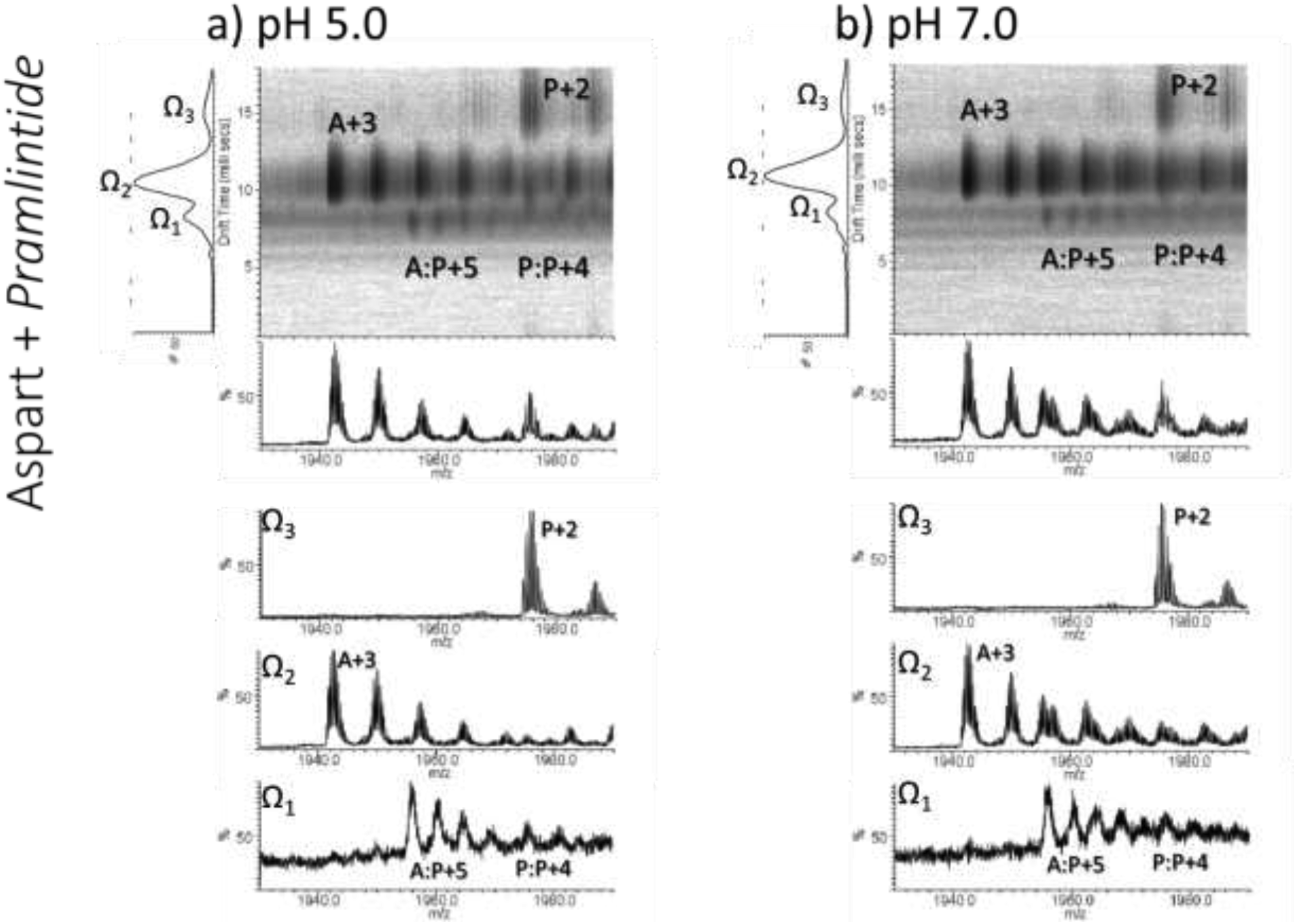
Distribution of oligomeric states of Aspart insulin and pramlintide accessed by electrospray ionization – ion mobility – mass spectrometry (ESI-IMS-MS). Aspart insulin (**A**) and pramlintide (**P**) were combined at 50 μM each in 50 mM ammonium acetate pH 5.0 (a) or pH 7.0 (b) and assayed for ESI-IMS-MS. Monomers are represented by a single letter (either **A** or **P**) and homodimers or hetero-dimers are identified by the combination of their corresponding letters. Their respective charge state, represented by number following the ‘plus’ sign, is shown adjacent to the identification of the oligomeric state. The ion intensity in the drift time (*d*t) axis x *m/z* is plotted in square root scale, and in linear scale for the total ESI-MS spectrum along the whole *d*t axis shown below the *d*t x *m/z* panel. A stripping of the *d*t x *m/z* space comprising specific *d*t regions (Ω_1_, Ω_2_ and Ω_3_) as assigned in the figure are shown for clarity.

**Figure 3.**
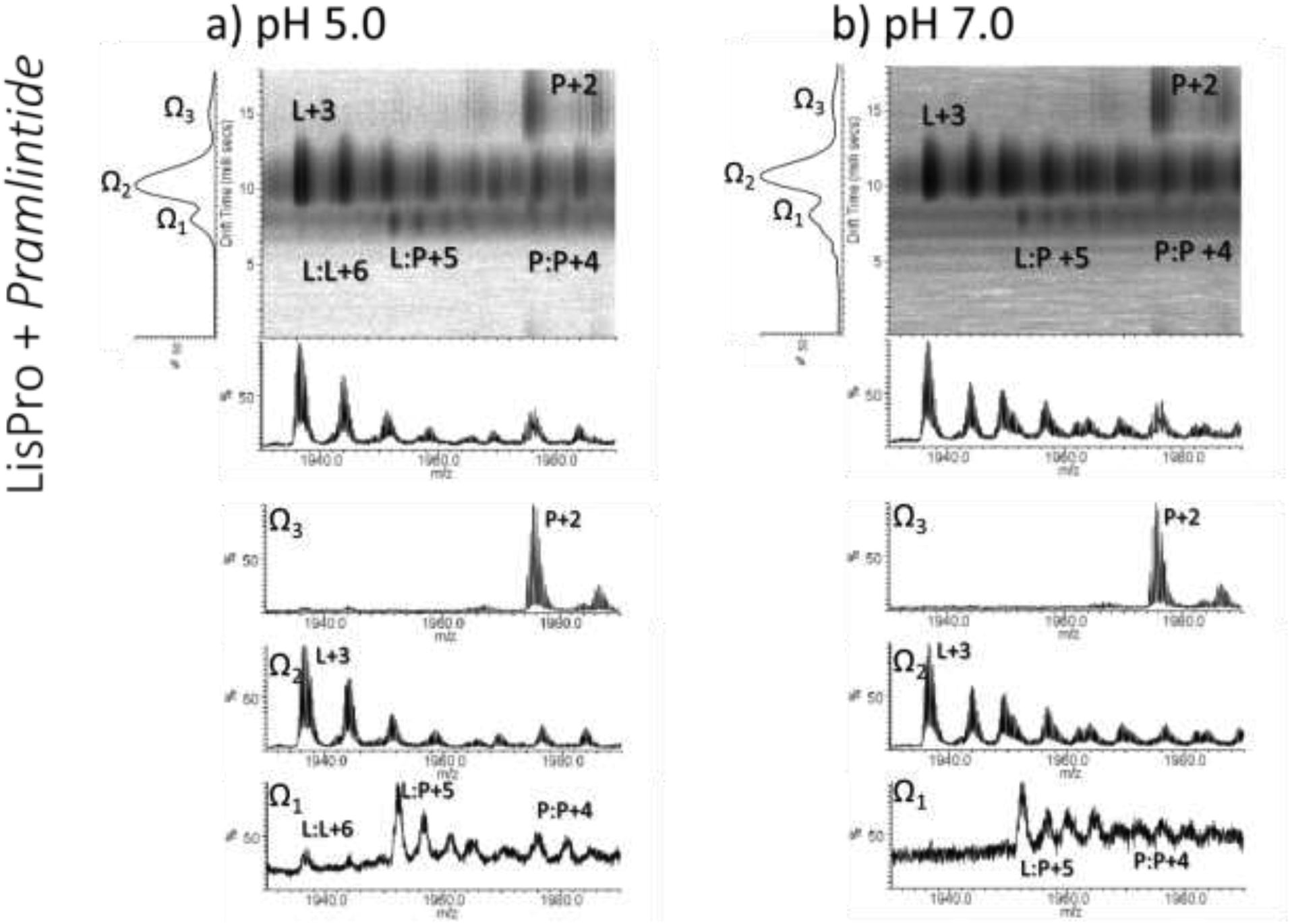
Distribution of oligomeric states of LisPro insulin and pramlintide accessed by electrospray ionization – ion mobility – mass spectrometry (ESI-IMS-MS). LisPro insulin (**L**) and pramlintide (**P**) were combined at 50 μM each in 50 mM ammonium acetate pH 5.0 (a) or pH 7.0 (b) and assayed for ESI-IMS-MS. Monomers are represented by a single letter (either **L** or **P**) and homodimers or hetero-dimers are identified by the combination of their corresponding letters. Their respective charge state, represented by number following the ‘plus’ sign, is shown adjacent to the identification of the oligomeric state. The ion intensity in the drift time (*d*t) axis x *m/z* is plotted in square root scale, and in linear scale for the total ESI-MS spectrum along the whole *d*t axis shown below the *d*t x *m/z* panel. A stripping of the *d*t x *m/z* space comprising specific *d*t regions (Ω_1_, Ω_2_ and Ω_3_) as assigned in the figure are shown for clarity.

**Figure 4.**
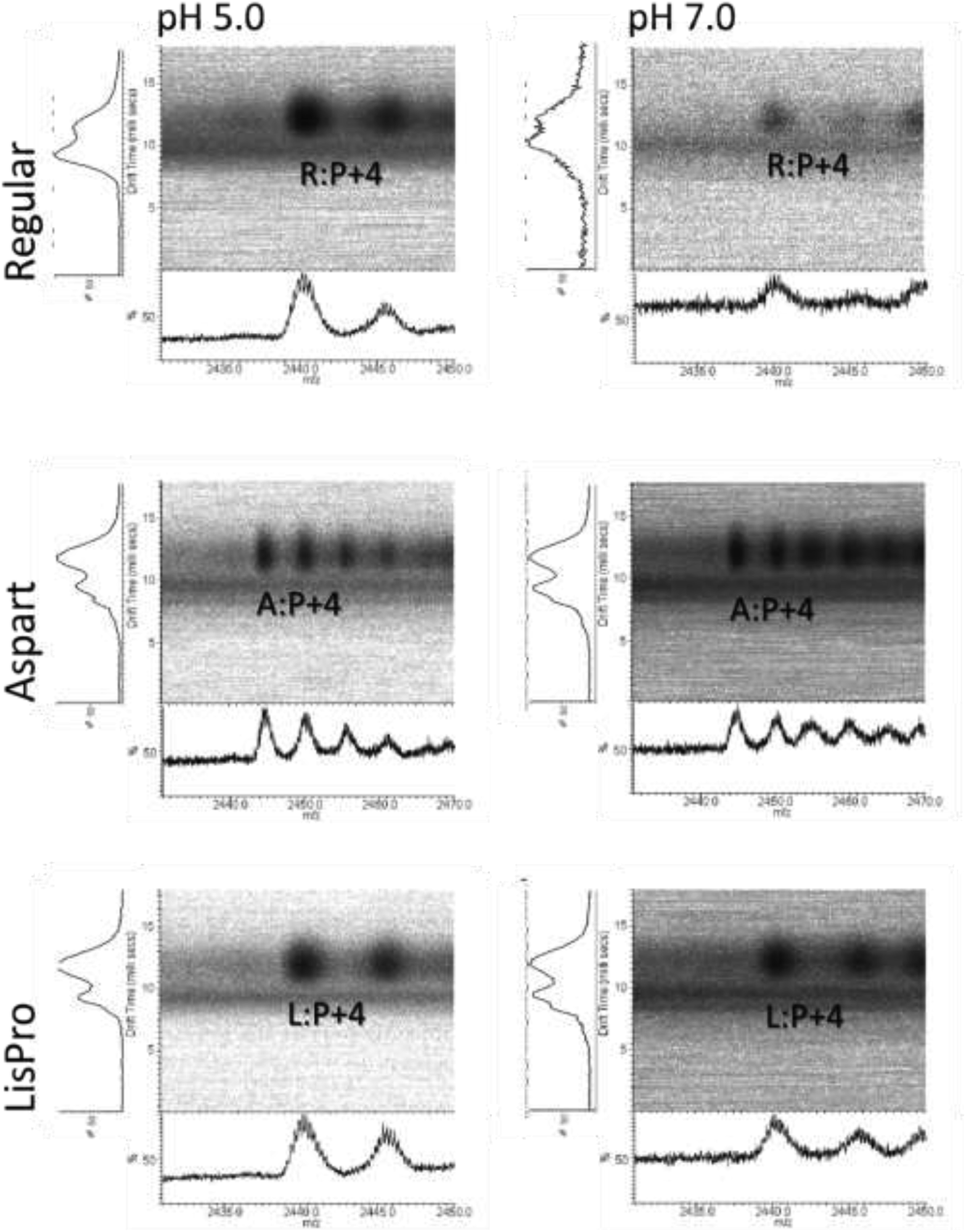
Identification of the +4 heterodimers of insulin and pramlintide by electrospray ionization – ion mobility – mass spectrometry (ESI-IMS-MS). Insulin (**R = regular** or **A = Aspart** or **L = LisPro**) and pramlintide (**P**) were combined at 50 μM each in 50 mM ammonium acetate pH 5.0 or pH 7.0 and assayed for ESI-IMS-MS. The ion intensity in the drift time (*d*t) axis x *m/z* is plotted **in square root scale, and in linear scale** for the total ESI-MS spectrum along the whole *d*t axis shown below the *d*t x *m/z* panel.

### Aggregation kinetics of insulin and pramlintide

– The drug substances insulin (42–44) and pramlintide (17) are known for their inherent propensity to association into amyloid aggregates under particular circumstances, although the drug medicines formulated with them are known for their physico-chemical stability at their final formulated and recommended storage conditions.

We thus decided to evaluate the propensity for amyloid aggregation of both pramlintide and insulin, separated and as combined preparation at 25 °C at pH 7.0 (representative of the pH of insulin formulations; **Table 1**), and at an acidic milieu. However, insulin undergoes acid-induced deamidation at its carboxy terminus, and it is catalyzed intramolecularly by protonated Asn^A21^. Increasing pH can decrease deamidation rate, being more stable pH 5.0 (45), which coincides with the beginning of acid-stabilization of insulin (46). Given these chemical stability issues (**Table 1**), we decided to test pH 5.0.

**Table 1.** Composition of commercial formulations of insulin variants and pramlintide.

|  | **Concentration** | **pH** | **Buffer** | **Other components** |
| --- | --- | --- | --- | --- |
| **<sup>1</sup>Regular Insulin** | 100 IU/mL<br>(aprox 3.45 mg/mL) | 7.0 – 7.8 | - | NaOH/HCl<br>16 mg/mL Glycerin<br>2.5 mg/mL *m*-cresol |
| **<sup>2</sup>Aspart Insulin** | 100 IU/mL<br>(aprox 3.45 mg/mL) | 7.2-7.6 | 1.25 mg Na<sub>2</sub>HPO<sub>4</sub> | NaOH/HCl<br>16 mg/mL Glycerin<br>1.50 mg/mL Phenol<br>1.72 mg/mL *m*-cresol<br>0.58 g/mL NaCl |
| **<sup>3</sup>LisPro Insulin** | 100 IU/mL<br>(aprox 3.45 mg/mL) | 7.0 – 7.8 | 1.88 mg Na<sub>2</sub>HPO<sub>4</sub> | NaOH/HCl<br>16 mg/mL Glycerin<br>3.15 mg/mL *m*-cresol |
| **<sup>4</sup>Pramlintide (vial)** | 600 µg/mL | 4.0 | 32 mM AcONa | 43 mg/mL d-mannitol<br>2.25 mg/mL *m*-cresol<br>1.53 mg/mL Acetic Acid<br>0.61 mg/mL Sodium Acetate |

From leaflet in https://www.accessdata.fda.gov/
1. Humulin R® or Novolin R® 2. NovoLog® 3. Humalog® 4. Symlin® - https://www.fda.gov/ohrms/dockets/ac/01/briefing/3761b1_05_Pharmacology.htm

An amyloid aggregation assay was conducted using 50 μM each polypeptide in 50 mM buffer pH 7.0 or pH 5.0, and ThT as amyloid fluorescent probe. Pramlintide showed a time-dependent increasing ThT fluorescence compatible with the formation of amyloid material at pH 7.0 (17), being more prominent in sodium phosphate (**Fig. 5A**) than in ammonium acetate (**Fig. 5B**) or sodium acetate (**Fig. 5C**) solutions. Instead, pramlintide showed no detectable increase in ThT fluorescence in these buffers at pH 5.0 (**Fig. 5D, 5E** and **5F**), confirming its enhanced stability at acidic milieu regardless of the buffer agent.

**Figure 5.**
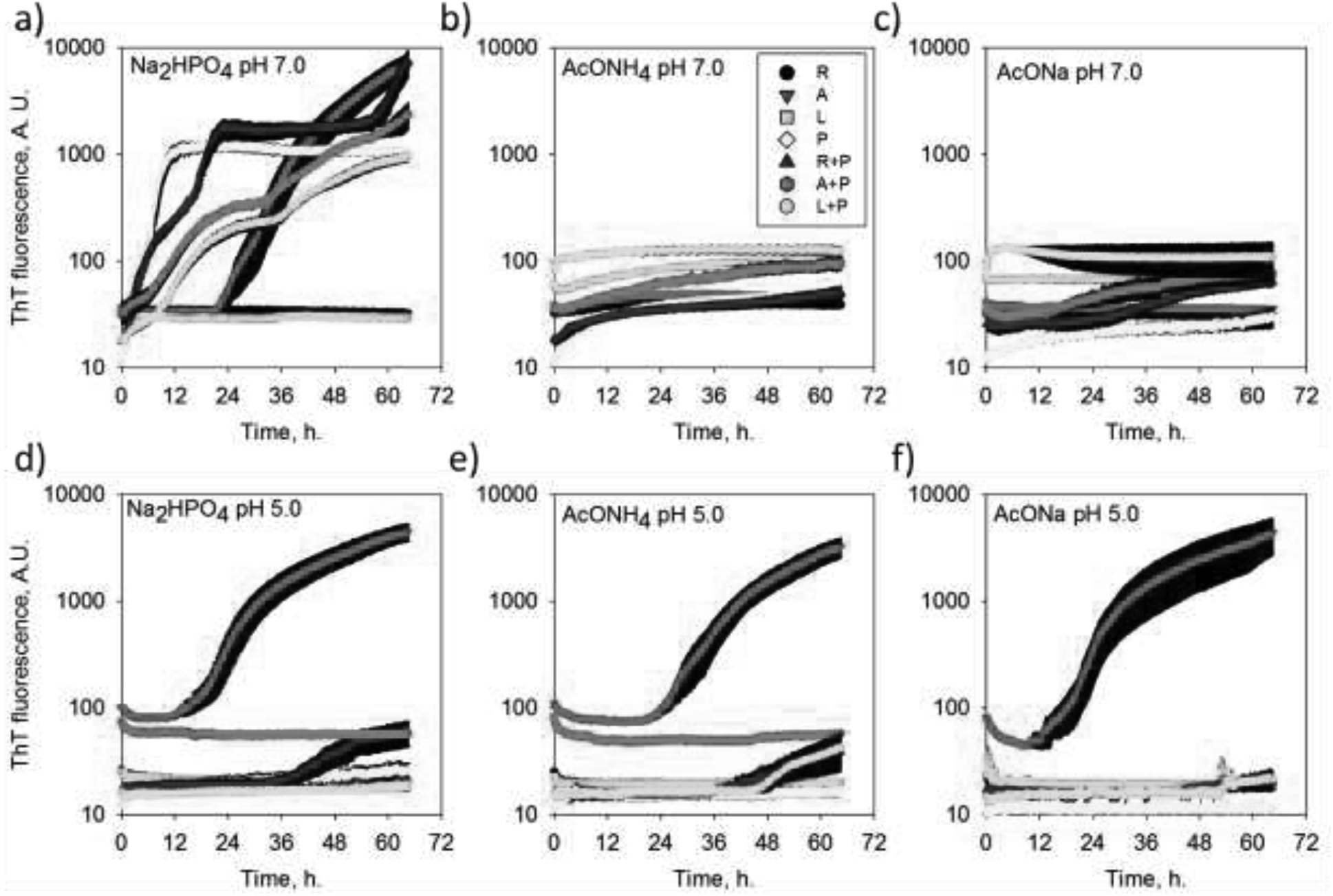
Aggregation kinetics of pramlintide and insulin analogues. Pramlintide and insulin analogues (regular, Aspart and LisPro, 50 μM each) were tested for amyloid aggregation at 25 °C alone or in combination, in 50 mM buffer sodium phosphate (A and D), ammonium acetate (B and E) or sodium acetate (C and F) at pH 7.0 (A, B, C) and pH 5.0 (D, E and F). ThT (20 μM) was used as probe for amyloid aggregates.

Regular insulin behaved stable for at least up to 65 h at 25 °C at pH 7.0 (**Fig. 5A, 5B** and **5C**) and pH 5.0 (**Fig. 5D, 5E** and **5F**). Combination of regular insulin with pramlintide did not hampered the amyloid aggregation process, only a delay in aggregation kinetic in a few hours in phosphate pH 7.0 plateauing at the same level of pramlintide alone, followed by an increase in ThT fluorescence after about 68 h (**Fig. 5A**). In fact, while regular insulin and pramlintide behave stable when incubated separately for over 65 h in sodium phosphate (**Fig. 5D**) or ammonium acetate (**Fig. 5E**) buffers pH 5.0 at 25 °C, combination of both result in an increase in ThT fluorescence indicating lack of compatibility of this formulation at these assay conditions.

The Aspart insulin showed signs of amyloid aggregation after a lag phase of about 12 h at pH 5.0 in sodium phosphate (**Fig. 5D**), ammonium acetate (**Fig. 5E**) and sodium acetate (**Fig. 5F**). At pH 7.0 in sodium phosphate, the Aspart insulin also showed an extensive increase in ThT (**Fig. 5A**), and a less prominent increase in ThT at this pH in the presence of ammonium and sodium acetate salts (**Fig. 5B** and **5C**, respectively).

Aggregation of Aspart insulin at pH 7.0 in phosphate may seem unusual given that it is formulated in the presence of this buffering agent. However, this aggregation assay requires diluting the protein 8 fold from its original formulation at about 600 μM (100 IU/mL) down to 50 μM, which may prompt for dissociation and amyloid aggregation. Furthermore, the other formulation components such as glycerol and phenolic compounds are diluted in the same extend, which may result in further decreased stability. But the inherent property of the Aspart insulin is likely to be deterministic, given that the LisPro insulin is formulated at similar conditions of the Aspart insulin (**Table 1**), but show no increase in ThT fluorescence in phosphate pH 7.0 during the assay (**Fig. 5A**). Instead, LisPro insulin showed an increase in ThT fluorescence at pH 7.0 in the presence of acetate (**Fig. 5B, 5C**), with no considerable increase in ThT at pH 5.0 in the presence of either buffers (**Fig. 5D, 5E, 5F**). In conjunction, these data suggest that the Aspart is a less stable insulin analogue.

At pH 7.0, incubation of pramlintide with insulin, either regular, Aspart or LisPro, resulted in increase in ThT fluorescence (**Fig. 5**) compatible with the observation of amyloid fibrils, both in phosphate and in acetate solutions (**Fig. 6**). At pH 5.0, regular and Aspart insulin showed an increase in ThT fluorescence in the presence of sodium phosphate (**Fig. 5D**) or ammonium acetate (**Fig. 5E**), indicative of amyloid aggregation, with limited detection when in sodium acetate (**Fig. 5F**). Insulin LisPro, by its turn, was the most stable preparation alone or in the presence of pramlintide, in both sodium phosphate (**Fig. 5D**) and acetate (**Fig. 5F**).

**Figure 6.**
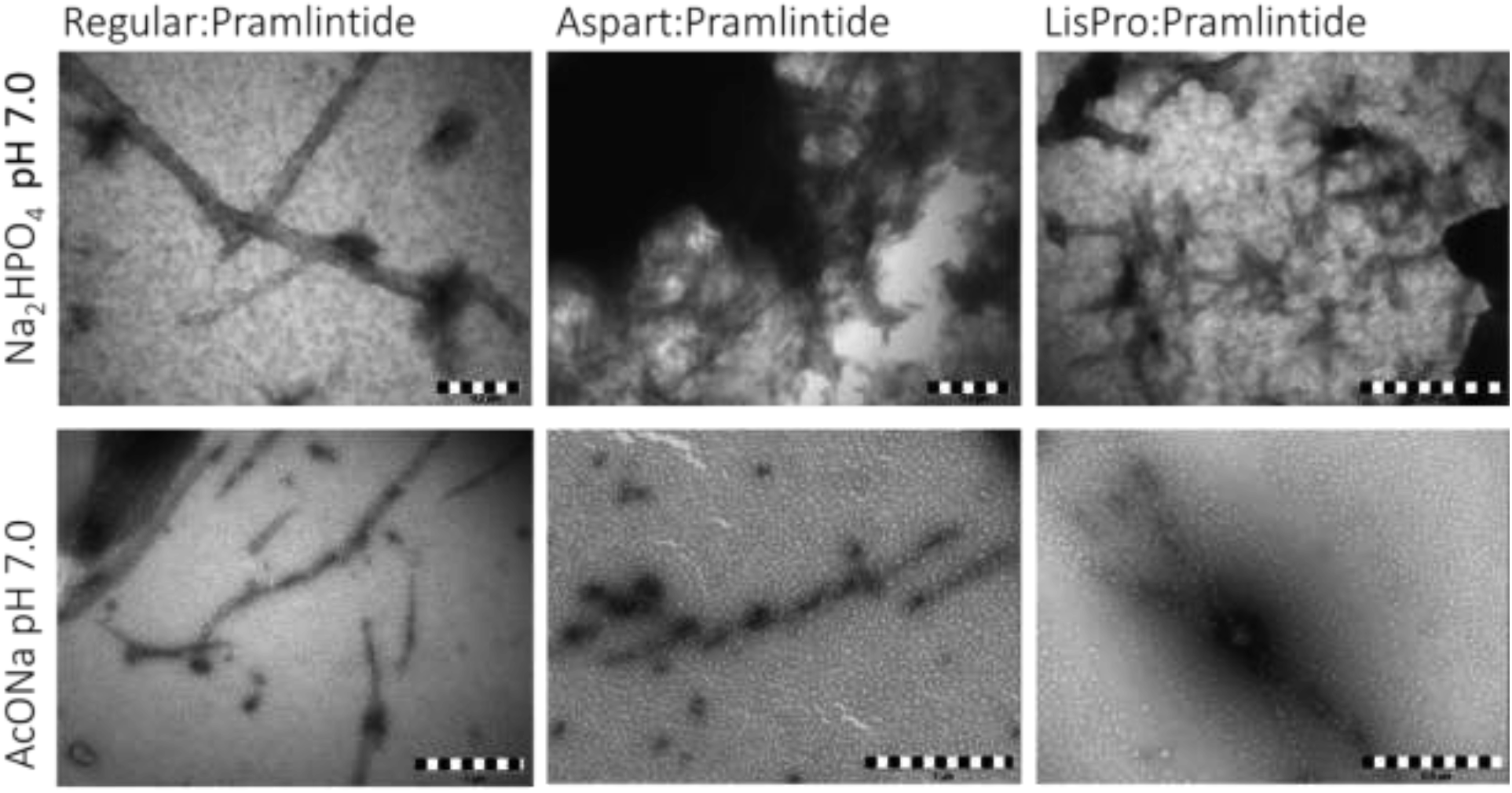
Morphologic characterization of amyloid aggregates by transmission electron microscopy (TEM). A combination of insulin:pramlintide at 50 μM each was allowed to aggregate for 3 days at 25 °C in 50 mM buffer and the material fixed onto a 300 mesh formvar-coated cupper grid and observed by TEM. Buffer, pH and insulin variants are according to indication.

These results indicated that both pH 7.0 and phosphate solutions are variables that may favor amyloid aggregation. Instead, sodium acetate is a compatible pharmaceutical ingredient and likely to provide the most favorable scenario in stabilization of the co-formulation of pramlintide at pH 5.0 with insulin, either regular, Aspart or Lispro. Under a therapeutic perspective, the co-formulation of fast-acting insulin with pramlintide would allow mimicking the hormonal behavior found in non-diabetic individuals, giving that regular insulin has a delayed onset, peak and duration of action compared to fast-acting insulin analogues (47). We have further tested the effect of *m*-cresol on the aggregation and no major issues were observed compared to control assay in the absence of the phenolic compound (**Fig. 7A**).

**Figure 7.**
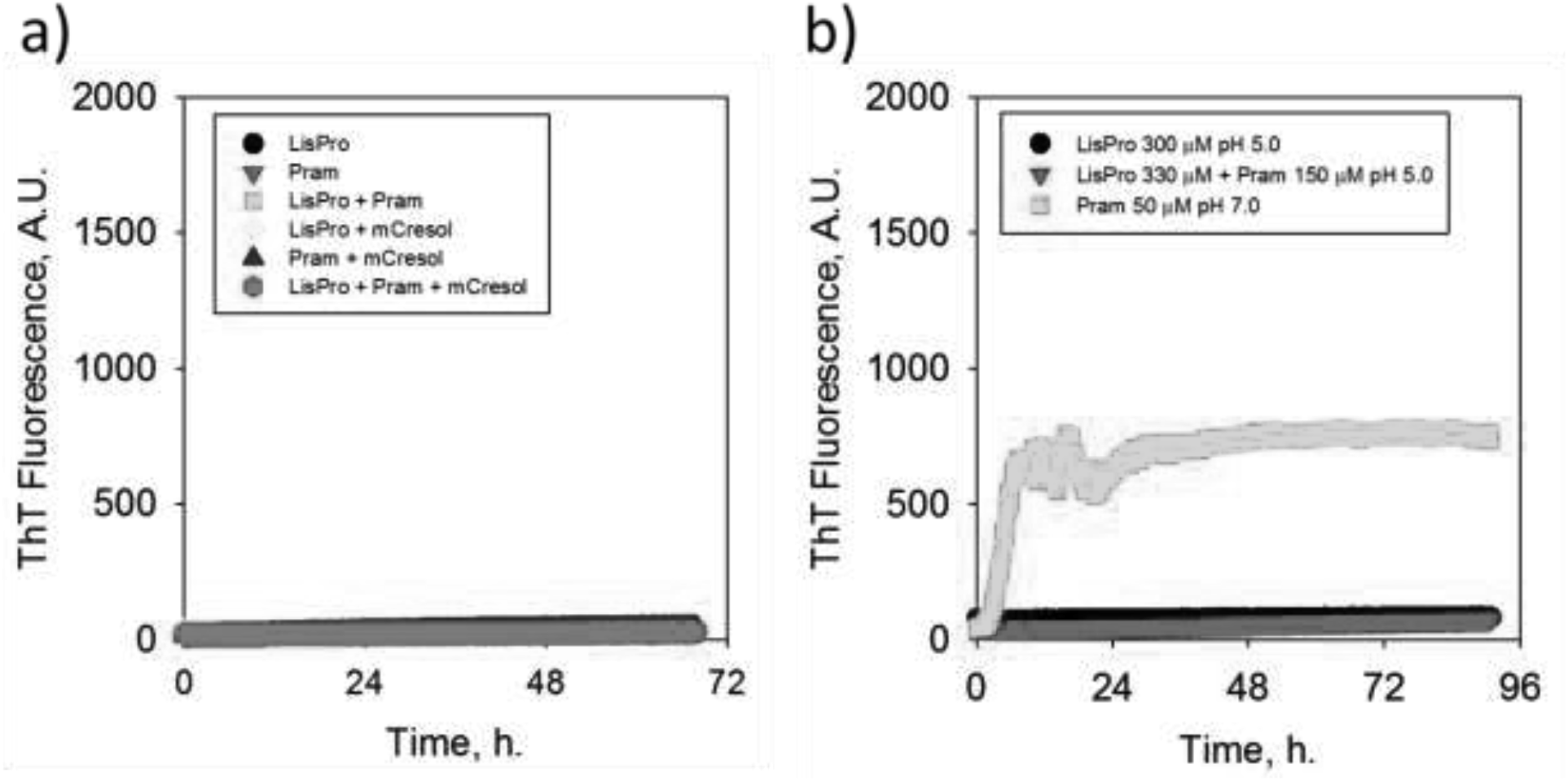
Stability of LisPro:Pramlintide co-formulation. a) Effect of m-cresol (2 mg/mL) on the aggregation profile of pramlintide and LisPro insulin (50 μM peptides), at *m*-cresol, 50 mM sodium acetate pH 5.0, 25 °C. **b)** Stability of co-formulated pramlintide and LisPro insulin at high concentration (300 μM) in 50 mM sodium acetate pH 5.0 at 25 °C. Pramlintide at 50 μM in 50 mM Na_2_HPO_4_ pH 7.0 is shown as positive aggregation control.

Insulin and pramlintide are both formulated at higher concentration than 50 μM. LisPro insulin is available at 100 IU/mL, about 3.5 mg/mL, which corresponds to 600 μM, while pramlintide is formulated at 600 μg/mL (vial), corresponding to 150 μM. Upon the introduction of pramlintide to a therapeutic scheme, an initial reduction in 50 % of insulin dose is recommended (26) with possible further adjustment. Based on these information, we decided to test the stability of a co-formulation of higher concentration of LisPro insulin (300 μM, about 50 IU/mL) and pramlintide (150 μM) in 50 mM sodium acetate pH 5.0. While the control pramlintide at low (50 μM) shows its typical fast amyloid aggregation (less than 12 h), the concentrated LisPro with and without pramlintide for over 72 h at 25 °C, as indicated by ThT (**Fig. 7B**). These data suggests the enhanced stability of the co-formulation of LisPro with pramlintide, both at high concentration at pH 5.0.

## DISCUSSION

Insulin is a vital therapeutic agent for T1DM, working as an adjunctive biopharmaceutical in advanced stages of T2DM. Amylin analogues are considered key player in the restoration of the physiology, due to its large spectrum of effects, as expected given their cosecretion by the pancreatic β-cells. The limitation so far in the use of pramlintide has been driven by its incompatibility upon co-formulation with insulin, preferably with fast-acting insulin in order to achieve similar PK profile. Separated administrations of insulin and amylin at distinct time before meals, varying doses and multiple injections sites acts as drawn-backs to the known therapeutic benefits of pramlintide, and thus a fixed ratio, single injection of a co-formulated insulin:pramlintide preparation would be desirable in mimicking physiological patterns (27).

We have shown here that both fast-acting insulin analogues Aspart and LisPro becomes stable in sodium acetate solution pH 5.0 while they become prone to amyloid aggregation at other assays conditions such as in phosphate solution and higher pH (**Fig. 5**). Attempts to co-formulate Aspart insulin with pramlintide failed during attempts to reduce the pH of the insulin. In opposition, the LisPro insulin behaved stable during this process allowing successful achievement of a combined, high-concentration co-formulation of this insulin analogue with pramlintide.

LisPro insulin is a fast-acting analogue, and that could be envisioned to be used both as a penfil and in automatic pump. The effectiveness of pramlintide in reduction of postprandial glucose excursions in combination with regular or LisPro insulin has already been demonstrated for both T1DM (48) and T2DM (49).

While the effective regular insulin is viewed as the more affordable therapeutic option compared to fast-acting insulin analogues (47), upon patent expiration for LisPro insulin combined formulation of this analogue with pramlintide could be considered for varying devices, vial-syringe, prefilled pen injectors or hybrid closed-loop insulin-pramlintide delivery system (“artificial pancreas”, such as in Clinical Trial NCT02814123 (50)).

This work has strengths such as reporting the formation of amyloid material with pramlintide alone at pH 7.0 or in combination with insulin within a few hours upon mixing, requiring cautionary actions concerning their use in combination, the potential effect of plastic interfaces (such as syringes and tubing) and room temperature (about 25 °C). Moreover, this work shows for the first time the compatibility of a fast-acting insulin with pramlintide, directing for the development of a co-formulated product, without the need for co-formulative copolymers such as oligosaccharides (51–53). Further studies are aimed to address long-term stability, a larger set of additional potential variables (54,55) such as adjuvant (glycerol, d-mannitol, phenolic compounds), as well as extending the range of investigated pH, temperature, and interfaces.

## ACKNOWLEDGMENTS

We would like to thanks Mr. Augusto Vieira and Dr. Dario Kalume for excellent support. We would like to thanks UEMP-UFRJ, CENABIO-UFRJ and INMETRO and their staff for the use of their analytical facilities. This research was supported by the Coordenação de Aperfeiçoamento de Pessoal de Nível Superior (CAPES), Conselho Nacional de Desenvolvimento Científico e Tecnológico (CNPq), Fundação de Amparo à Pesquisa do Estado do Rio de Janeiro Carlos Chagas Filho (FAPERJ). The funding agencies had no role in the study design, data collection and analysis, or decision to publish or prepare of the manuscript.

